# MaGplotR: a software for the analysis and visualization of multiple MaGeCK screen datasets through aggregation

**DOI:** 10.1101/2023.01.12.523725

**Authors:** Alejandro Matía, Maria M. Lorenzo, Duo Peng

**Affiliations:** Department of Biotechnology, Instituto Nacional de Investigación y Tecnología Agraria y Alimentaria – Consejo Superior de Investigaciones Científicas (INIA-CSIC), Madrid, Spain; Chan Zuckerberg Biohub, San Francisco, CA, USA

## Abstract

**Motivation:** MaGplotR analyzes multiple CRISPR screen datasets, identifies common hits, couples the hits to enrichment analysis and cluster plots, and produces publication-level plots. Output plots give information on the quality control of the screen data (e.g., sgRNA distribution) and show the best hits by aggregation from multiple screen experiments. To maximize comparability, rank is used to identify common hits. MaGplotR can also be used to analyze experiments where a control condition is used for multiple treatment conditions. MaGplotR is easy to use, with even just one argument.

**Availability and implementation:** MaGplotR is open-source code under MIT license. It is available at https://github.com/alematia/MaGplotR.

## Introduction

Genetic screens, and especially CRISPR/Cas9 screens are a revolutionary tool in the field of genomics. Under different selection conditions, the effect of single gene expression by knockout, knockdown or activation of expression can be analyzed and selected, obtaining differentially enriched or depleted hits. A list of bioinformatic tools have been created for the analysis of the screen results (sequencing data), with different statistical approaches to obtain a final list of hits [1–5]. Among these, MaGeCK (Model-based Analysis of Genome-wide CRISPR/Cas9 Knockout), is the most widely used and offers a rank list of hits based on their log2-fold change and p-value, using a modified robust ranking aggregation algorithm (RRA) [1].

Several bioinformatic tools directed to the visualization of the screen results have also been developed, such as MaGeCK Vispr and CRISPRAnalyzeR [6, 7]. To the best of our knowledge, no tools have been developed to analyze common hits from MaGeCK results of multiple experiments/conditions. Here, we introduce MaGplotR, a software for the analysis and visualization of multiple CRISPR screening datasets, including the option to use a control condition for baseline comparison. Specifically, MaGplotR is focused on the detection of the best common hits from multiple datasets by using the gene RRA ranking score instead of enrichment scores as the sorting criteria, which makes different datasets more comparable to each other, regardless the specific enrichment/depletion scores distribution of each experiment. The performance of the hits in the control experiment is visually enhanced with a specific plot to simplify the analysis of potential false positive hits. Also, MaGplotR uses the gene summary files from the MaGeCK test command and does not rely on the fastq or read counts files, simplifying the aggregation analysis for non-experienced users, and complementing the existing software tools.

### Software description

MaGplotR is a software written in R coding language. It can be executed from a Linux, macOS, or Windows command line. MaGplotR has a few dependencies (R packages) that can be installed either from within an R terminal, or from the command line using the installation script provided for non-experienced users [8–14]. The main program (MaGplotR.R), as well as the installation script, documentation, and example files are publicly available online at https://github.com/alematia/MaGplotR. For the analysis and generation of output plots and files from 4 screen experiments, the complete workflow (executing all optional arguments) can take less than 1 min on a standard computer.

### MaGplotR detects the best hits from aggregation of multiple datasets

MaGplotR takes one mandatory input: a folder with a collection of gene_summary.txt files generated from the MaGeCK RRA algorithm, each file representing an experimental condition or a different dataset. In each instance, MaGplotR generates up to 6 different plots and 2 intermediate data files. The first plots are the boxplot and the heatmap-an example is shown in Figure 1A-. For the boxplot, the log-fold change of each gene is represented as well as the box and whiskers of each experiment. Outliers are also represented to give a quick view of the distribution of the potential hits. Moreover, if a control experiment is used, the boxplot will include the control data. Comparing the scattering of LFC (against the control) can help to inform the magnitude of the selective pressure applied to the screens. A heatmap is plotted to compare the LFC of top hit genes across condition/datasets. The heatmap is sorted (row-wise) based on the average rank of each gene in the set of experiments, which aggregates the common hits from the different conditions/datasets. The color represents the LFC of each gene in the corresponding experiment and the number inside the cell corresponds to the rank of the gene. The number of hits shown in the heatmap can be chosen as an argument in the command line. Also, a horizontal dot plot will be generated complementary to the heatmap plot if a control file is supplied, easing the analysis of the performance of the potential hits in the control experiment, which can help to discard false positives-i.e. if a hit shows a enrichment behavior in the control of a positive enrichment experiment- (Fig 1B). We include a plot to examine the expression of top hit genes in a set of most commonly used human cell lines, based on data from The Human Protein Atlas [15] (Fig 1C). During the program execution, messages will be displayed indicating each step of the computing process.

**Fig 1.**
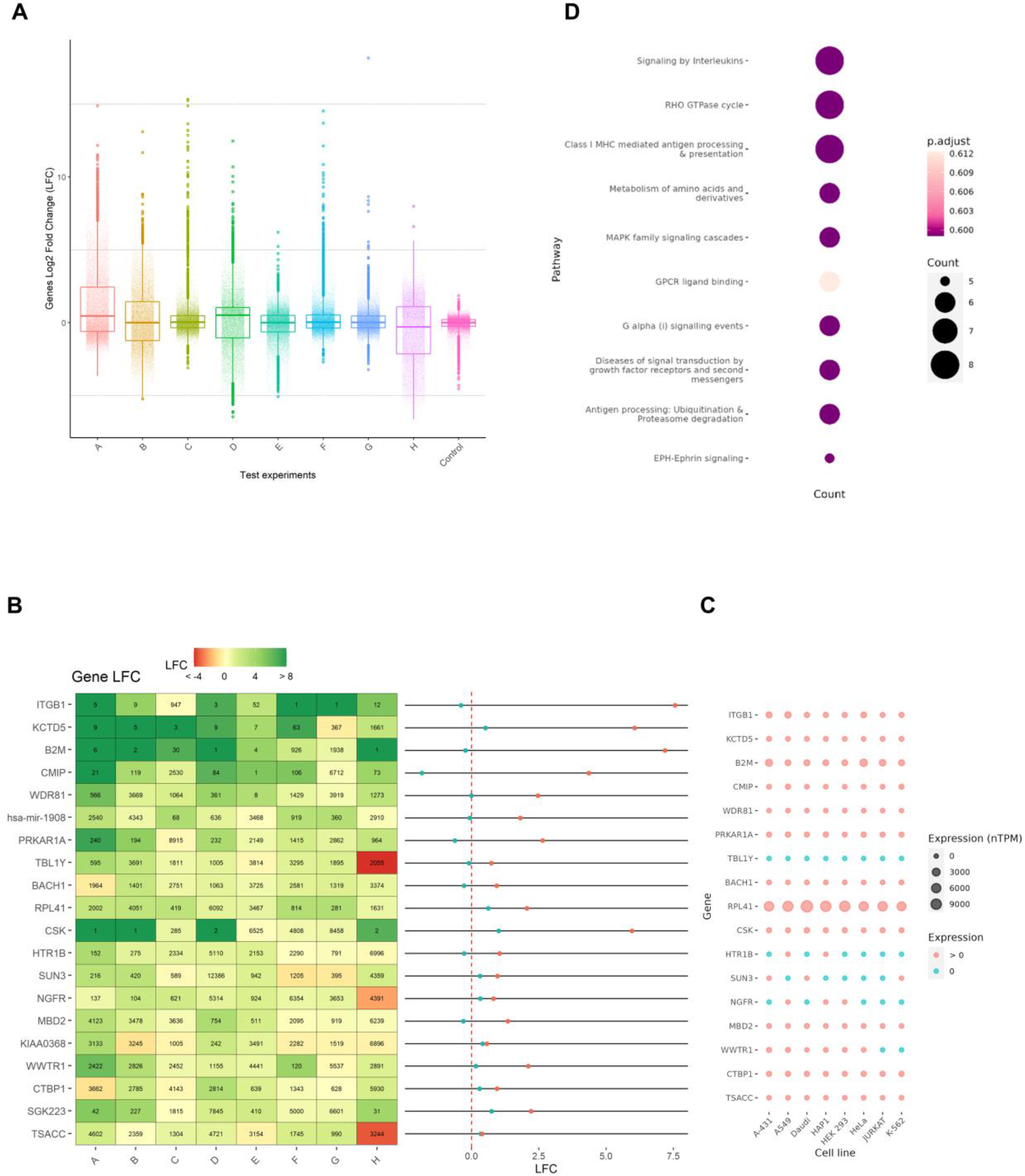
A. Boxplot representing the LFC distribution for each screen experiment. B. Heatmap and control dotplot. Heatmap showing the best aggregation hits. The colors indicate the LFC of each gene in each experiment. The number inside the cell is the rank of that particular gene in an experiment. The control dotplot is only generated in the case of using a control as option. C. Expression plot. Dotplot representing the number of transcripts per million of the top hits for the most representative cell lines. D. Pathway plot. Dotplot showing the number of genes assigned to a certain pathway.

### MaGplotR performs enrichment analysis and generates network cluster plots

MaGplotR includes a downstream analysis and visualization of the top 1 % of hits (by default, and configurable by the user) using Reactome Pathway Analysis and clusterProfiler, allowing the user to explore the enriched interactome with different degrees of tightness (Fig 1D). A list of the genes that belong to the most enriched pathway will be printed out. Also, Gene Ontology analysis (Biological Process, Cellular Component, Molecular Function) can be done optionally.

All plots but the cluster plot (pdf only) can be output in different formats (png, pdf, ps, jpeg, tiff, bmp).

### Utility and prospects

This tool has been tested using multiple datasets to offer robust and wide applicability [16–20]. Of note, this tool can be used to detect hits that were previously unnoticed, by aggregating common hits from different experiments, as is the case of GATA6 proviral gene for SARS-CoV-2, which we identified as the 4^th^ hit using rearranged Biering et al. datasets [19], and has been confirmed by Israeli et al. [21].

## Conclusion

MaGplotR is a fast, lightweight, and easy-to-use tool for elegant visualization that outputs several unique plots to facilitate the analysis of multiple screen results. Specifically developed to improve analysis through visualization of data from several experiments, we expect it to be useful for the community of researchers.

## Funding

A.M. was recipient of a predoctoral contract from Subprograma Estatal de Formación, Programa Estatal de Promoción del Talento y su Empleabilidad en I+D+I, Spain.

## Notes

https://github.com/alematia/MaGplotR

### Competing Interest Statement

The authors have declared no competing interest.

## References

1. Li, W., et al., MAGeCK enables robust identification of essential genes from genome-scale CRISPR/Cas9 knockout screens. Genome Biol, 2014. 15(12): p. 554.

2. Diaz, A.A., et al., HiTSelect: a comprehensive tool for high-complexity-pooled screen analysis. Nucleic Acids Res, 2015. 43(3): p. e16.

3. Hart, T. and J. Moffat, BAGEL: a computational framework for identifying essential genes from pooled library screens. BMC Bioinformatics, 2016. 17: p. 164.

4. Allen, F., et al., JACKS: joint analysis of CRISPR/Cas9 knockout screens. Genome Res, 2019. 29(3): p. 464–471.

5. Yu, J., J. Silva, and A. Califano, ScreenBEAM: a novel meta-analysis algorithm for functional genomics screens via Bayesian hierarchical modeling. Bioinformatics, 2016. 32(2): p. 260–7.

6. Li, W., et al., Quality control, modeling, and visualization of CRISPR screens with MAGeCK-VISPR. Genome Biol, 2015. 16: p. 281.

7. Cui, Y., et al., VISPR-online: a web-based interactive tool to visualize CRISPR screening experiments. BMC Bioinformatics, 2021. 22(1): p. 344.

8. Davis, T.L., Optparse: Command Line Option Parser. CRAN: the Comprehensive R Archive Network. 2018.

9. Hadley, W., Reshaping Data with the reshape Package. Journal of Statistical Software, 2007. 21(12): p. 1–20.

10. Hadley, W., et al., Welcome to the tidyverse. Journal of Open Source Software, 2019. 4(43): p. 1686.

11. Carlson, M., org.Hs.eg.db: Genome wide annotation for Human. R package version 3.8.2. 2019.

12. Yu, G. and Q.Y. He, ReactomePA: an R/Bioconductor package for reactome pathway analysis and visualization. Mol Biosyst, 2016. 12(2): p. 477–9.

13. Wu, T., et al., clusterProfiler 4.0: A universal enrichment tool for interpreting omics data. Innovation (Camb), 2021. 2(3): p. 100141.

14. Bengtsson., H., et al., Package ‘matrixStats’. 2022.

15. Karlsson, M., et al., A single-cell type transcriptomics map of human tissues. Sci Adv, 2021. 7(31).

16. Dukhovny, A., et al., A CRISPR Activation Screen Identifies Genes That Protect against Zika Virus Infection. J Virol, 2019. 93(16).

17. Wenfang S. Tan, et al., Genome-wide CRISPR knockout screen reveals membrane tethering complexes EARP and GARP important for Bovine Herpes Virus Type 1 replication. BioRxiv, 2020.

18. Emily, H., et al., A modular CRISPR screen identifies individual and combination pathways contributing to HIV-1 latency. BioRxiv, 2022.

19. Biering, S.B., et al., Genome-wide bidirectional CRISPR screens identify mucins as host factors modulating SARS-CoV-2 infection. Nat Genet, 2022. 54(8): p. 1078–1089.

20. Matia, A., et al., Identification of beta2 microglobulin, the product of B2M gene, as a Host Factor for Vaccinia Virus Infection by Genome-Wide CRISPR genetic screens. PLoS Pathog, 2022. 18(12): p. e1010800.

21. Israeli, M., et al., Genome-wide CRISPR screens identify GATA6 as a proviral host factor for SARS-CoV-2 via modulation of ACE2. Nat Commun, 2022. 13(1): p. 2237.

